# Data-Driven Symbolic Higher-Order Epistasis Discovery with Kolmogorov-Arnold Networks

**DOI:** 10.1101/2025.10.31.685894

**Authors:** Oankar R. Patil, Kamran Shazand, Benoit Marteau, Yuxuan Shen, May D. Wang

## Abstract

Many human diseases are polygenic conditions that arise from a complex interplay of interactions between multiple genes at different loci, but currently most Genome-Wide Association Studies (GWAS) largely only consider the main additive effects of single nucleotide polymorphisms (SNPs), resulting in a missing heritability problem in some complex traits. Identifying non-additive interactions, or epistasis, at a higher-order could aid in filling this gap, but it is computationally difficult due to the massive search space involved. Current epistasis detection approaches struggle with non-cartesian higher order interactions and lack inherent explainability. We present a novel deep learning (DL) approach, EPIstasis Discovery with Kolmogorov-Arnold Networks (EPIK), a data-driven, modular, and symbolically representable framework. We also introduce a novel approach for higher-order XOR (a non-Cartesian type) interaction detection, utilized in EPIK’s XOR detection module. EPIK slightly outperforms other DL approaches on simulated pure epistasis interactions benchmark in average F1 score. It outperforms other, general, traditional epistasis detection approaches on simulated mixed epistasis detection datasets and real-world GWAS datasets of Arabidopsis Thaliana. Finally, EPIK recovers a known gene interaction between MAPT and WNT3 for Parkinson’s Disease (PD) while also suggesting a more complex interaction between MAPT, WNT3, and another gene, KANSL1.

## 1 Introduction

Many human diseases, including Parkinson’s disease, are polygenic and shaped by interactions between multiple genes across loci, known as epistasis [7, 28]. Epistasis is a key candidate for the “missing heritability” problem in complex traits, yet most Genome-Wide Association Studies (GWAS) focus only on additive effects of single nucleotide polymorphisms (SNPs) [19]. Detecting higherorder, non-additive interactions is computationally challenging: for *n* SNPs, evaluating all possible interactions up to order *m* requires 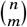 combinations, an *o* (*n*^*m*^) operation. The high dimensionality of GWAS data further compounds this issue by invoking the curse of dimensionality [6].

Recent work has pursued non-exhaustive search strategies, particularly deep learning (DL) methods, for efficient higher-order epistasis detection. These include multilayer perceptrons (MLPs) with biological priors, layer-wise relevance propagation (e.g., DeepCOMBI) [26] [17], convolutional and transformer-based models [14] [10], and relief-based algorithms [8]. While promising, these approaches remain black-box models and struggle with non-Cartesian interactions such as XOR, which can model more relevant interactions than additive or multiplicative models in some cases [2].

Kolmogorov-Arnold Networks (KANs) offer a potential solution. Unlike MLPs with fixed activations, KANs use learnable B-spline functions on edges, enabling compact parameterization, improved generalization, and intrinsic explainability [16]. Their design is inspired by the Kolmogorov-Arnold representation theorem, which decomposes multivariate functions into sums of univariate functions, aligning well with sparse compositional structures such as true causal SNPs embedded within noisy features. Importantly, KANs yield interpretable symbolic or graphical representations of detected interactions.

We propose EPIstasis discovery with Kolmogorov-Arnold Networks (EPIK), a modular, data-driven framework for uncovering higher-order epistasis from SNP datasets with intrinsic symbolic explainability. We evaluate EPIK on simulated epistasis and real GWAS datasets, comparing it against state-of-the-art (SOTA) methods, and apply it to Parkinson’s disease. EPIK successfully recovers a known MAPT–WNT3 interaction in Parkinson’s Disease (PD) and suggests a novel three-way interaction that includes another gene, KANSL1.

Our contributions are as follows:

- A novel polynomial-time approach for efficient higherorder (≥ 4) XOR epistasis detection, improving performance on non-Cartesian interactions.
- To the best of our knowledge, this is the first application of KANs to epistasis discovery, providing interpretable symbolic and graphical representations while outperforming or matching other non intrinsically-interpretable approaches.

## 2 Methodology

### 2.1 Datasets

#### 2.1.1 Simulated Pure Epistasis Datasets

We utilized a publicly available collection of datasets for higher-order pure epistasis detection, utilizing the synthetic test datasets seen in [10], which were generated utilizing PyToxo [9] for penetrance table generation and GAMETES [25] for simulating interacting SNP combinations that satisfy Minor Allele Frequency (MAF), heritability, number of SNPs, sample size, control vs case ratio parameters. Such simulated data produced falls under pure, strict epistasis models, being worst-case scenarios for disease association due to only being observable if all k-loci are included. There were a total of 25 datasets to evaluate performance in a wide variety of k-way epistasis detection scenarios, covering Additive, Multiplicative, Thresholding, XOR interactions.

#### 2.1.2 Simulated Realistic Mixed Epistasis Datasets

To evaluate performance on mixed epistasis interaction types/realistic genetic noise, we used the “msprime” population genetic simulator [3] and simulated the genetic ancestries of 1,600 individuals (800 AFR, 400 EUR, 400 ASN) with recombination/mutations. We implemented penetrance tables for two mixed epistasis scenarios and adding biallelic mutation, generating two datasets with varying MAFs, each containing ∼93,500 SNPS. The first contained 2-way Multiplicative (MAF: 0.17, 0.41), 2-way Recessive (MAF: 0.12, 0.31), 2-way XOR (MAF: 0.25, 0.23) with 780 Cases/820 Controls. The second contained 3-way Multiplicative (MAF: 0.05, 0.19, 0.18) and 5-way XOR (MAF: 0.31, 0.37, 0.41, 0.45, 0.40) with 757 Cases/843 Controls.

#### 2.1.3 Arabidopsis Thaliana GWAS datasets

We utilized GWAS data of Arabidopsis Thaliana [1] for additional experiments with traditional non-DL epistasis approaches [27] [8] [13] [22] [18]. The datasets consisted of SNPs (∼24,500) across all of its 5 chromosomes, aiming to identify genes involved in resistance of various bacterial effector phenotypes (avrRPM1, avrPphB).

#### 2.1.4 Parkinson’s Disease Dataset

For deeper insight into PD which has recently been heavily investigated for epistasis interactions [5] [21], we utilized genomic patient samples from the UKBioBank for SNPs across chromosome 17 [4]. Specifically, we selected 1790 patients afflicted with a PD diagnosis based on patient samples with ICD-10 codes of G20. We then selected 18159 controls patients, thus following a 10:1 control:case ratio. For quality control of our patient control population, we followed the same procedure as previous meta-analysis studies for epistasis in PD GWAS [5].

### 2.2 Architecture Overview

EPIK overall consists of four epistasis interaction type detection modules that each propose a candidate set of SNPs involved in epistasis interaction before being fed to their respective KANs. Each KAN attempts to learn the interaction from the candidate SNPs, prune SNPs it finds unhelpful, and produce a symbolic representation of the interaction. Our approach operates on the usage of machine learning for intrinsic explainability and modularity, aiming to work with algorithmic improvements as the field evolves.

### 2.3 Additive Detection Module

For higher order additive detection, we utilize inspiration from approaches already established in literature that follow heuristic search and information gain for epistasis detection [23] [24]. Our approach works by doing a stepwise floating search strategy that combines spline-based nonlinear modeling with Bayesian Information Criterion (BIC) optimization [11]. We pre-screen SNPs by computing univariate BIC scores using spline-transformed genotypes to select top candidates, then perform iterative forward-backward search: adding SNPs that minimize BIC when their genotype sum is transformed via spline basis functions (capturing nonlinear interactions) and removing previously added SNPs if exclusion improves BIC. This approach models additive joint effects through the sum of genotypes while controlling complexity via BIC’s penalty term.

### 2.4 XOR Detection Module

We present a novel approach for detecting multi-way XOR-type epistatic interactions in high-dimensional genotype-phenotype data. Due to the nature of an XOR interaction, no single SNP/subset of SNPs is guaranteed a large marginal correlation with the phenotype, but only when all causal SNPs are in combination with one another. As such, any algorithm for this task has two major goals:

- Filtering the initial high-dimensional dataset to a reduced subset of candidate SNPs that retain the causal SNPs.
- Recognizing combinations of SNPs as an XOR interaction that has high correlation with the phenotype.

Before addressing the problem of a high-dimensional dataset, we first consider what would be a relevant and useful “mechanism” for detecting such an XOR interaction if we were somehow given the actual set of causal SNPs. If we take a moment to consider what an XOR interaction actually shows up as, we can see from the definition that it can be boiled down to the idea of, “If an odd number of the inputs are true, the output will be true”.

Thus, the problem transforms into: if given some set, is the parity of this set even or odd? As such, we need some sort of approach that can compute the parity of a given set. The mechanism we use to detect this parity pattern is computing the Walsh Coefficient of a given set.

For a set of SNP indices *S* = *i*_1_, *i*_2_, …, *i*_*m*_, the Walsh coefficient is:

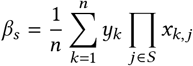

Here:

- *x*_*k,j*_ ∈ +1, −1 is the coded genotype of individual k at SNP
- The product 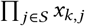 is +1 if an even number of carriers in *S* are present, and –1 if an odd number are present.

We take the absolute value of *β*_*s*_. Now that we know we are trying to maximize the Walsh coefficient given various sets, the question becomes how can we effectively select those sets, especially given a high-dimensional dataset. If we were to try to compute the Walsh coefficients for all subsets of *p* SNPs, there would be 2^*p*^ subsets, which would be computationally intractable in a high-dimensional setting. Thus we now lead into discussion on how to address the first goal of effectively filtering SNPs and choosing relevant subsets of SNPs to find the true XOR interaction. We consider whether some sort of correlation based approach would be a cheaper yet effective initial screen.

Based on the correlation derivation we did, shown in detail in the Appendix, we can see that for a *k*-way XOR with independent SNPs, with *p* being MAF:

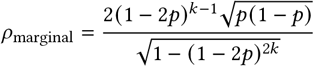

This effectively denotes the following relationship:

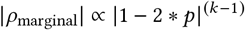

Thus, we see that as MAF approaches 0.5, this term -> 0, and quickly as *k* grows (higher order). We plot this relationship in **Figure 2**.

**Figure 1.**
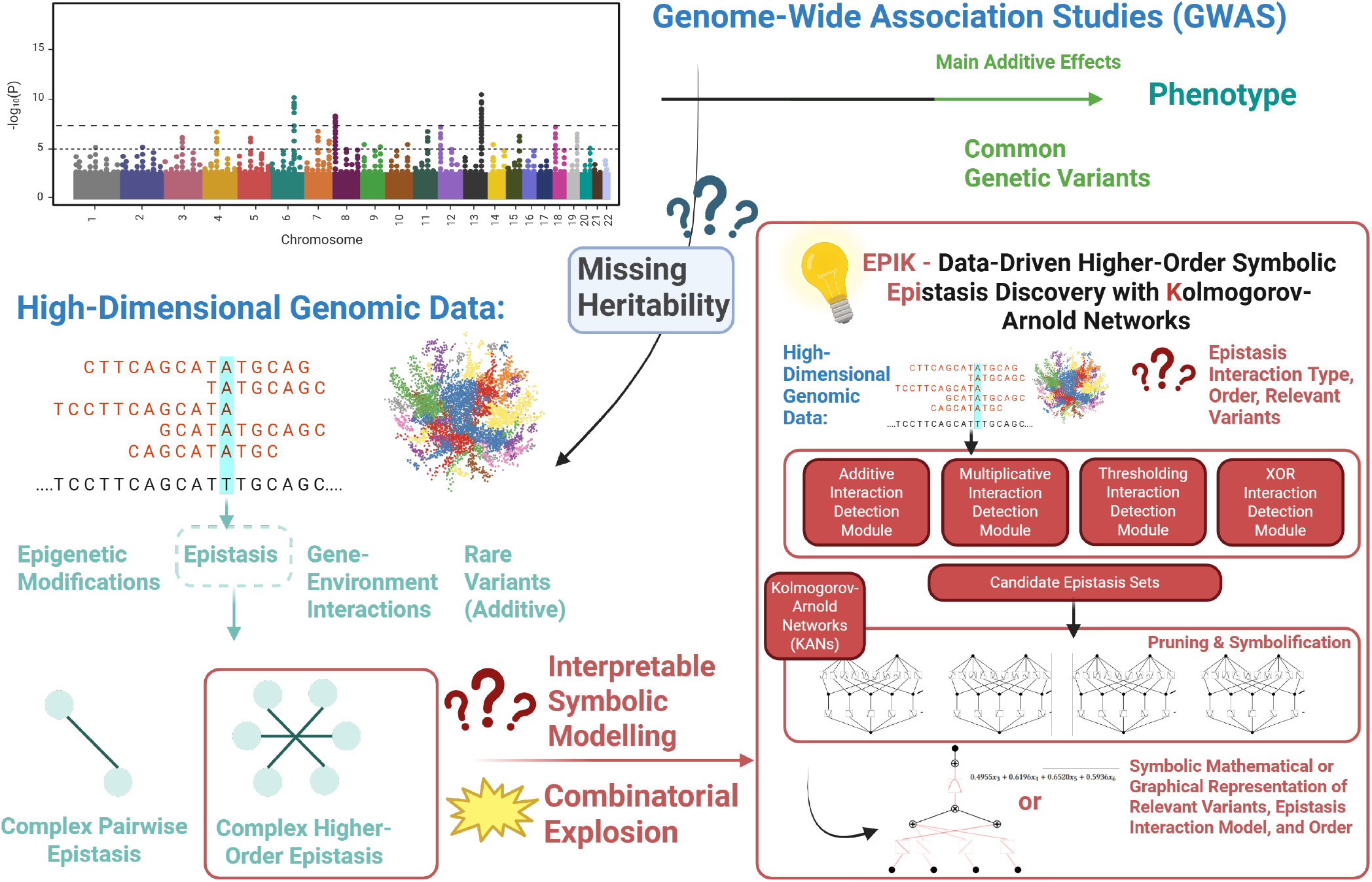
The missing heritability problem in GWAS may stem from factors such as epistasis, gene–environment interactions, and rare variants. Within epistasis, higher-order interactions pose major computational and interpretability challenges. We propose EPIK, a framework combining optimized higher-order detection methods with Kolmogorov–Arnold Networks for efficient and interpretable epistasis discovery.

**Figure 2.**
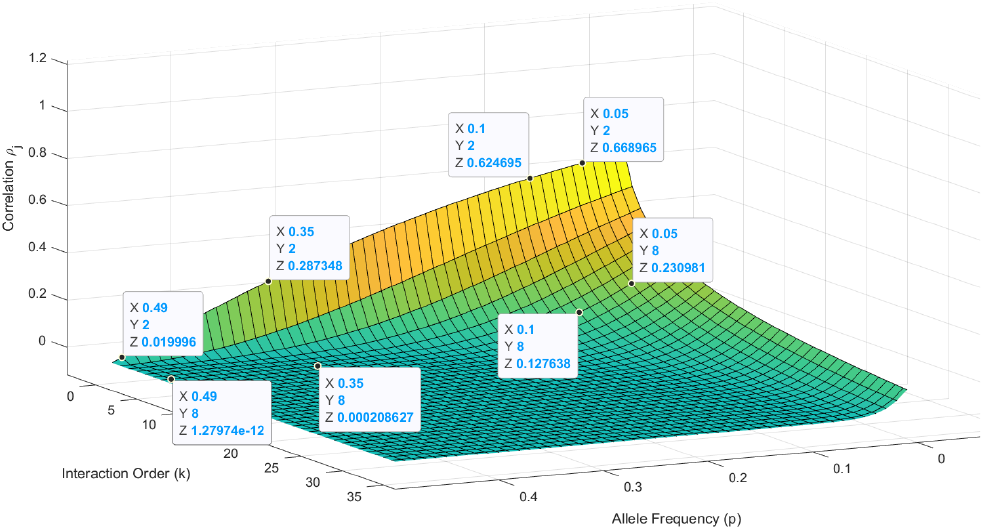
We plot the region of final k-way XOR correlation for given MAF and interaction order k. We also plot specific cases for second order interactions and eighth order interactions at varying MAF values.

As such, we consider each SNP with random sets of SNPs at a given order and compute the parity-transformed set *k*-way correlation with the phenotype. We then rank each SNP as candidates based on how well their sets generally correlated with the phenotype and take a user-defined number of initial candidates, hoping to capture small “spikes” in initial correlation. For high-dimensional datasets, scaling-wise we see this shifts us from the intractable 2^*p*^ to effectively *n* \**p*, where *n* is the number of random sets for each SNP. From these indvidual SNP candidates, we then do a beam search up to the user-defined max order, computing the Walsh coefficient for each potential set. The final output is the set with the highest Walsh coefficient.

SNPs with low MAF values in an XOR interaction are thus more likely to be captured, but it is still possible to capture SNPs with higher MAF values but is just more difficult. This can be mitigated by increasing the number of initial candidates parameter. We further detail our approach in **Algorithm 1 and 2** in the Appendix.

### 2.5 Multiplicative Detection Module

We detect higher-order multiplicative epistasis through a parallelized beam search guided by log-likelihood ratio (LLR) tests, beginning with mutual information-based pre-screening of top candidate SNPs to reduce combinatorial complexity, similar to other approaches in literature for lower-order interaction detection [12].

### 2.6 Thresholding Detection Module

Our approach detects higher-order threshold epistasis through an adaptive beam search guided by point-biserial correlation optimization, beginning with pre-screening SNPs based on their marginal correlation with binarized phenotypes. Our approach evaluates candidate SNP sets by computing minor allele counts across loci and dynamically determining optimal activation threshold ≥*k* (within a proportional range) where the rule *counts* ≥ *k* maximizes absolute correlation with the phenotype. Using beam search with backtracking, it expands candidate sets order-by-order while retaining only top-performing combinations based on correlation strength.

### 2.7 KANs Hyperparameters, Pruning, Symbolification and Prediction

For each candidate set of SNPs, a KAN is trained to simultaneously learn which SNPs from the candidate set are most relevant, whether the initial order of the candidate set can be reduced, and a symbolic representation (mathematical or graphical) of how these SNPs interact together in consideration of their proposed epistasis interaction type. We take a quick aside to clarify such symbolic representations are an inherent property of KANs and not something achieved through another method such as how LRP or Neural Interaction Detection (NID) might be applied to MLPs. All of the KANs have an initial first layer width equivalent to an initial proposed candidate set for each interaction type. We provide the exact networks depth/widths and other parameters in the Appendix. All of the KANs discussed used auto-symbolification/pruning parameters of 0.7 for the r2 threshold and a simplicity weight of 0.5. We utilize a stratified 80/20 split to train and test each KAN.

We tuned each KAN for each epistasis interaction type considered in our framework. We did this by separately generating and extracting only the causal sets of SNPs for each interaction type using PyToxo and GAMETES, leaving the benchmark datasets untouched, and doing a hyperparameter search/different network sizes on these. The problem on how to optimally design KANs for different tasks is still open, but based on [16][15], we treat KANs like more interpretable MLPs and start as small as possible before first increasing width and then depth. We cap depth to a maximum of five, as [15] discusses how deep KANs roughly become an identity map in the later layers.

### 2.8 Experimental Design

Our first experiment evaluates the performance of EPIK on simulated pure and mixed epistasis interaction scenarios. We compare (F1 Score) against other DL approaches [10] [17] [14] on the 25 simulated pure epistasis interactions datasets benchmark described in **Section 2.1.1**. We also evaluate performance of EPIK against non deep-learning epistasis detection approaches [27] [8] [13] [22] on mixed epistasis interaction datasets described in **Section 2.1.2**.

Our second experiment evaluates the performance of EPIK with well-established epistasis detection approaches [27] [8] [13] [22] [18] on two real-world GWAS datasets described in **Section 2.1.3**. Finally, our last experiment applies EPIK for PD in chromosome 17 with a magnitude of 10^4^ input SNPs considered, using the dataset as described in **Section 2.1.4**. We first do traditional LD-pruning, with a *r*^2^ threshold of 0.7, a MAF filter of 0.01, and population covariate control through PCA before then feeding the dataset to EPIK. We evaluated EPIK on a single Nvidia A100 GPU with 4 CPU cores and 16 GB memory. We utilized up to 4 A100 GPUs as needed for the other approaches, dependent on their computational needs.

## 3 Results

### 3.1 Simulated Epistasis Detection

We evaluate the performance of EPIK against SOTA DL methods on the simulated datasets described in **Section 2.1.1**. We present performance against the current SOTA, a distributed transformer approach [10] with the remaining DL approaches in the Appendix. For the pure epistasis interaction types, we find that EPIK outperforms or is on par with the other SOTA DL method by average F1 score in the majority of *k*-way scenarios, shown in **Table 1a**. We also present the performance of EPIK against non deep-learning approaches on simulated mixed epistasis datasets described in **Section 2.1.2**, where EPIK outperforms the other approaches in number of causal SNPs found, shown in **Table 1**’s mixed epistasis sub-tables **Table 1b** and **Table 1c**.

**Table 1:** Simulated Epistasis Interaction Results.

**(a) Pure Epistasis Interactions (F1 Score)**
| (Pure Interaction Type) DL Approach | 2-way | 3-way | 4-way | 5-way | 6-way | 7-way | 8-way |
| --- | --- | --- | --- | --- | --- | --- | --- |
| (Add) Distributed Transformer | 0.92 | 0.78 | 0.89 | 0.97 | 0.99 | 0.98 | 0.81 |
| (Add) EPIK | 1.0 | 0.75 | 0.89 | 0.91 | 0.92 | 0.88 | 0.89 |
| (XOR) Distributed Transformer | 0.44 | 0.35 | 0.26 | 0.26 | 0.21 | 0.14 | 0.22 |
| (XOR) EPIK | 1.0 | 0.86 | 1.0 | 0.91 | 0.92 | 0.93 | 0.94 |
| (Mult) Distributed Transformer | 0.57 | 0.73 | 0.72 | 0.79 |  |  |  |
| (Mult) EPIK | 0.8 | 0.67 | 0.73 | 0.72 |  |  |  |
| (Thr) Distributed Transformer | 0.67 | 0.67 | 0.58 | 0.44 | 0.54 | 0.58 | 0.62 |
| (Thr) EPIK | 1.0 | 0.67 | 0.73 | 0.67 | 0.72 | 0.57 | 0.57 |

**(b) Mixed Epistasis Interactions — Scenario 1**
|  | EPIK | Multi SURF | Multi SURF* | Fast LMM | PLINK BOOST | w-test |
| --- | --- | --- | --- | --- | --- | --- |
| Total Number of Causal SNPs Found (Detected/Total) | 4/6 | 2/6 | 1/6 | 1/6 | 2/6 | 3/6 |
| 2-way XOR SNPs (Detected/Total) | 2/2 | 1/2 | 1/2 | 0/2 | 0/2 | 2/2 |
| 2-way Mult SNPs (Detected/Total) | 1/2 | 1/2 | 0/2 | 1/2 | 2/2 | 0/2 |
| 2-way Recessive SNPs (Detected/Total) | 1/2 | 0/2 | 0/2 | 0/2 | 0/2 | 1/2 |

**(c) Mixed Epistasis Interactions — Scenario 2**
|  | EPIK | Multi SURF | Multi SURF* | Fast LMM | PLINK BOOST | w-test |
| --- | --- | --- | --- | --- | --- | --- |
| Total Number of Causal SNPs Found (Detected/Total) | 4/8 | 2/8 | 1/8 | 1/8 | 2/8 | 2/8 |
| 5-way XOR SNPs (Detected/Total) | 3/5 | 1/5 | 1/5 | 0/5 | 0/5 | 1/5 |
| 3-way Mult SNPs (Detected/Total) | 1/3 | 1/3 | 0/3 | 1/3 | 1/3 | 1/3 |

**Table 2:**
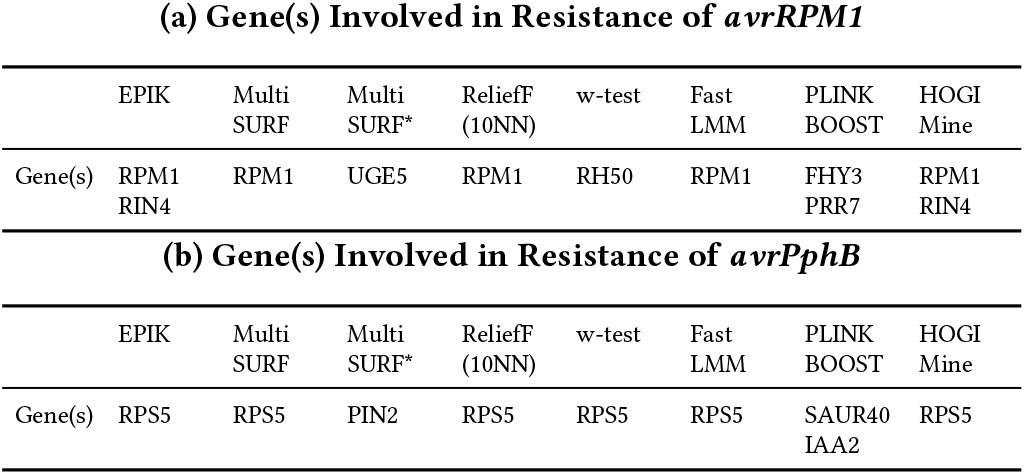
*A. Thaliana* GWAS Experiments.

**(a) Gene(s) Involved in Resistance of *avrRPM1***
|  | EPIK | Multi SURF | Multi SURF* | ReliefF (10NN) | w-test | Fast LMM | PLINK BOOST | HOGI Mine |
| --- | --- | --- | --- | --- | --- | --- | --- | --- |
| Gene(s) | RPM1<br>RIN4 | RPM1 | UGE5 | RPM1 | RH50 | RPM1 | FHY3<br>PRR7 | RPM1<br>RIN4 |

**(b) Gene(s) Involved in Resistance of *avrPphB***
|  | EPIK | Multi SURF | Multi SURF* | ReliefF (10NN) | w-test | Fast LMM | PLINK BOOST | HOGI Mine |
| --- | --- | --- | --- | --- | --- | --- | --- | --- |
| Gene(s) | RPS5 | RPS5 | PIN2 | RPS5 | RPS5 | RPS5 | SAUR40<br>IAA2 | RPS5 |

### 3.2 *A. Thaliana* GWAS Experiments Results

We evaluate EPIK against well-established epistasis detection approaches as described in **Section 2.1.1** in **Table 2**. Out of these approaches, HOGIMine is the only one that uses biological priors. For resistance of avrRPM1, EPIK and HOGIMine recovered RPM1/RIN4, while MultiSURF, RelifF-10NN, Fast-LMM recovered only RPM1, with RPM1/RIN4 being involved [29]. For resistance of avrPphB, EPIK, MultiSURF, RelifF-10NN, w-Test, Fast-LMM, HOGIMine recovered RPS5, with RPS5/PBS1 being involved [20].

### 3.3 Parkinson’s Disease Results

We show EPIK’s findings for PD on Chromosome 17 in comparison to prior literature (**Table 3**), a genome-wide non-exhaustive epistasis meta-analysis [5]. We show EPIK’s intrinsic symbolic representation, suggesting an XOR interaction (**Figure 3**).

**Table 3:** Parkinson’s Disease Epistasis Findings (Chr 17)

|  |  |
| --- | --- |
| EPIK’s Proposed SNPs (rsID) | rs2696439, rs7210219, rs199503 |
| EPIK’s Proposed Associated Genes | MAPT, WNT3, KANSL1 |
| Genes associated with significant Epistasis Interaction in Literature | MAPT, WNT3 |
| Significant Gene EQTL from MAPT-WNT3 in Literature | KANSL1, LRRC37A, LINC02210, KANSL1-AS1, ARL17B, LRRC37A2 |

**Figure 3.**
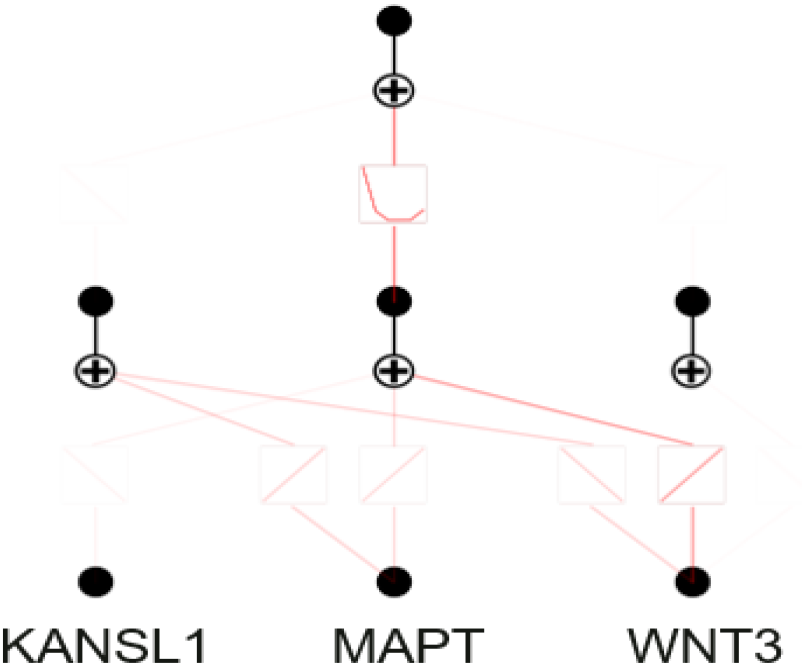
EPIK’s intrinsic symbolic representation suggests a potential 3-way XOR interaction between KANSL1, MAPT, and WNT3 for Parkinson’s Disease in Chromosome 17.

## 4 Discussion

Overall, across simulated datasets with varying epistasis interaction types and orders, EPIK matched or outperformed other DL and traditional methods in most scenarios. A key strength of EPIK is its intrinsic symbolic representation, which enables pruning of unlikely SNPs. For example, in the pure 7-way additive scenario (where EPIK underperformed against Distributed Transformer), we recovered the following symbolic expression:

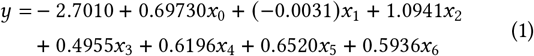

Here, *x*_1_ has a near-zero negative coefficient, correctly indicating it as a noisy non-epistasis SNP that was pruned.

Not all interactions yield concise mathematical expressions. In such cases, EPIK’s symbolic graphical representation highlights pruned SNPs, noisy candidates, and potential interactions. For instance, in the 3-way Pure Threshold Interaction (**Fig. 4**), one proposed SNP, highlighted in yellow, is faintly represented, signaling possible noise, which is consistent with the ground truth. The remaining SNPs in blue were correctly pruned and the ones in Red were the causal SNPs.

**Figure 4.**
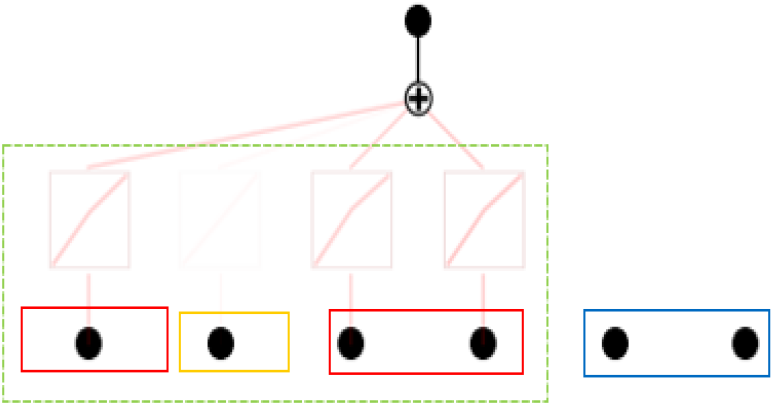
EPIK’s intrinsic symbolic graphical representation of the 3-way Threshold Interaction scenario.

Applying EPIK to chromosome 17 in Parkinson’s disease, we identified a 3-way XOR interaction between KANSL1, MAPT, and WNT3. The symbolic graph (**Fig. 3**) emphasized MAPT and WNT3 as the primary interaction, with KANSL1 faint, suggesting noise or a regulatory role. Literature confirms MAPT and WNT3 as a significant non-additive epistasis pair [5], while additional EQTL/*E*^2^*QT L* analysis implicates KANSL1 and nearby genes. Despite reduced sample size compared to prior studies, EPIK successfully recovered the main MAPT/WNT3 interaction.

We note that the XOR module lacks global statistical guarantees due to its non-exhaustive search, tested subsets yield exact correlation estimates, with detection power depending on effect size, beam width, and candidate selection. For the full framework, statistical guarantees can be incorporated via candidate–test splits, likelihood ratio tests, and BH-FDR correction. We provide a detailed discussion of these procedures and considerations for higher-order interactions in the Appendix.

EPIK’s XOR detection module design is biased in that it has a higher chance of finding SNPs in an interaction if they have lower MAF values, necessitating future work to mitigate this.

## 5 Conclusion

EPIK presents a symbolically representative approach to epistasis discovery. KANs are still a very nascent class of neural networks and thus optimal architecture design for this task is still greatly open for future work. Further work could also investigate integration with functional annotations and other biological prior knowledge like known pathways, chromatin accessibility, or regulatory annotations, into the interpretation or prioritization steps which could potentially greatly improve the ability to find biologically relevant and robust higher order epistasis interactions.

Overall, EPIK introduces a novel higher-order XOR epistasis detection algorithm and KAN-based approach for data-driven intrinsic symbolic discovery of higher-order epistasis, outperforming or matching other approaches on simulated and real GWAS datasets, and suggesting a more complex epistasis interaction in 17q21.31 for PD.

## Acknowledgments

This research was supported by Shriners Children’s Hospital and the Georgia Institute of Technology through the Early Onset Scoliosis (EOS) Project. This research was supported in part through research cyberinfrastructure resources and services, including the AI Makerspace of the College of Engineering, provided by the Partnership for an Advanced Computing Environment (PACE) at the Georgia Institute of Technology, Atlanta, Georgia, USA. This research has been conducted using the UK Biobank Resource under Application 17984.

## A Appendix

### A.1 Marginal Correlation for XOR Epistasis

Consider a *k*-way XOR model for the phenotype *Y* :

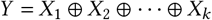

where *X*_*i*_ ∼ Bernoulli (*p*) are independent SNPs. The correlation *ρ* _*j*_ between SNP *X* _*j*_ and *Y* is:

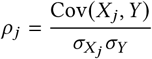

*Covariance Decomposition:*

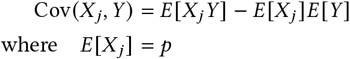

*Expectation of Y* : *Y* = 1 when an odd number of *X*_*i*_ = 1, being a sum that follows binomial distribution:

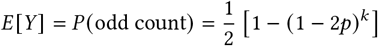

*Expectation of X* _*j*_*Y* : Conditional on *X* _*j*_ = 1:

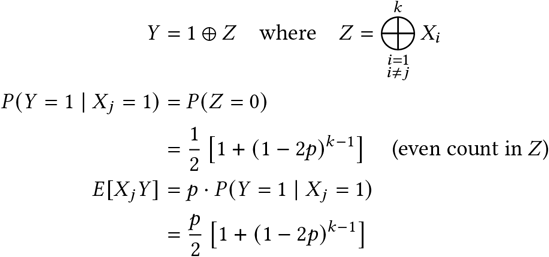

*Compute Covariance:*

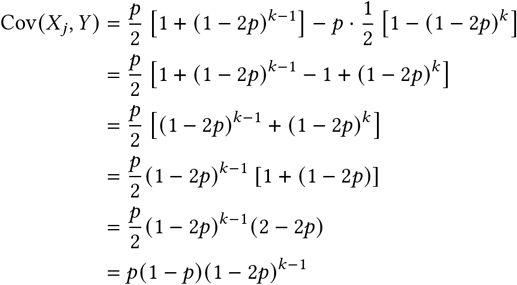

*Variance Components:*

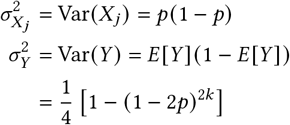

*Final Correlation:*

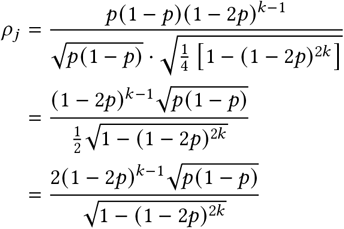

### A.2 XOR Detection Module - Algorithms

Shown in **Algorithm 1 and 2**.

### A.3 Simulated Pure Epistasis Interaction Results

We present the full comparison of EPIK against the DL approaches discussed in the main paper in **Table 4**. F1 score for EPIK was calculated by taking the union of final proposed candidate SNP sets from each KAN module after their pruning process.

**Table 4:** Simulated Epistasis Interaction Results (F1 Score)

| (a) Pure Additive |  |  |  |  |  |  |  |
| --- | --- | --- | --- | --- | --- | --- | --- |
| Approach | 2-way | 3-way | 4-way | 5-way | 6-way | 7-way | 8-way |
| Distributed Transformer | 0.92 | 0.78 | 0.89 | 0.97 | 0.99 | 0.98 | 0.81 |
| EPIK | 1.0 | 0.75 | 0.89 | 0.91 | 0.92 | 0.88 | 0.89 |
| DeepCombi | 0.47 | 0.29 | 0.43 | 0.57 | 0.46 | 0.26 | 0.41 |
| CNN | 0.34 | 0.32 | 0.54 | 0.42 | 0.39 | 0.51 | 0.35 |
| (b) Pure XOR |  |  |  |  |  |  |  |
| Approach | 2-way | 3-way | 4-way | 5-way | 6-way | 7-way | 8-way |
| Distributed Transformer | 0.44 | 0.35 | 0.26 | 0.26 | 0.21 | 0.14 | 0.22 |
| EPIK | 1.0 | 0.86 | 1.0 | 0.91 | 0.92 | 0.93 | 0.94 |
| DeepCombi | 0.13 | 0.06 | 0.02 | 0.00 | 0.01 | 0.00 | 0.00 |
| CNN | 0.07 | 0.01 | 0.00 | 0.01 | 0.00 | 0.00 | 0.00 |
| (c) Pure Multiplicative |  |  |  |  |  |  |  |
| Approach | 2-way | 3-way | 4-way | 5-way |  |  |  |
| Distributed Transformer | 0.57 | 0.73 | 0.72 | 0.79 |  |  |  |
| EPIK | 0.8 | 0.67 | 0.73 | 0.72 |  |  |  |
| DeepCombi | 0.09 | 0.26 | 0.21 | 0.12 |  |  |  |
| CNN | 0.24 | 0.23 | 0.03 | 0.00 |  |  |  |
| (d) Pure Threshold |  |  |  |  |  |  |  |
| Approach | 2-way | 3-way | 4-way | 5-way | 6-way | 7-way | 8-way |
| Distributed Transformer | 0.67 | 0.67 | 0.58 | 0.44 | 0.54 | 0.58 | 0.62 |
| EPIK | 1.0 | 0.67 | 0.73 | 0.67 | 0.72 | 0.57 | 0.57 |
| DeepCombi | 0.39 | 0.43 | 0.32 | 0.18 | 0.11 | 0.10 | 0.09 |
| CNN | 0.22 | 0.12 | 0.19 | 0.01 | 0.02 | 0.01 | 0.00 |

### A.4 Sensitivity Analysis of XOR Detection Module for Pure XOR under Moderate MAF

We performed our method on 3-way XOR datasets with moderate MAF (0.3) (due to computational generation limitations with GAMETES we did not go higher) and had the initial candidate subset parameter vary from [33, 32, 31, 30], representing a reduction to only 0.033% to 0.03% of the original high-dimensional SNP dataset, shown in **Table 5**. We see that the first causal SNP lost was the 32nd highest marginal correlation. Thus, we overall see the core mechanisms of the XOR modules performance lies in MAF values and initial candidate subset parameter.

**Table 5:**
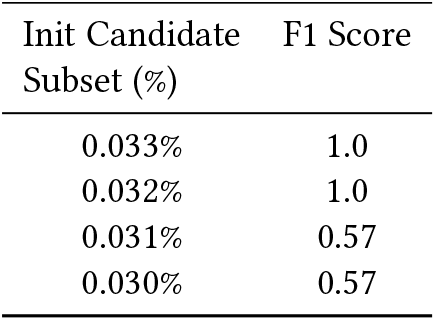
Moderate MAF impact on 3-Way XOR Detection.

### A.5 KAN Models Details

Additive KAN directly goes to a size [1] width output, number of grid intervals is set to 5, a piecewise polynomial order of 3, and the use of identity as the base function. XOR Kan has a width list of [20, 1] after the first layer, number of grid intervals is set to 5, a piecewise polynomial order of 3, and the use of Sigmoid Linear Unit (SiLU) as the base function. Multiplicative KAN has a width list of [[0, 1], 1], number of grid intervals set to 20, a piecewise polynomial order of 3, the use of identity as the base function, and a multiplication arity of either 2 or the closest scalar of half of the candidate set. Multiplicative KAN utilizes the recent multiplication node architecture developed for KANs [15]. Finally, the Thresholding KAN directly goes to a size [1] width output, number of grid intervals is set to 5, a piecewise polynomial order of 3, and the use of identity as the base function.

### A.6 Multiplicative KAN

We highlight EPIK’s usage of the multiplication node for KAN architectures, developed in [15], in its intrinsic symbolic graphical representations for Multiplicative epistasis scenarios in **Figure 5**.

**Figure 5.**
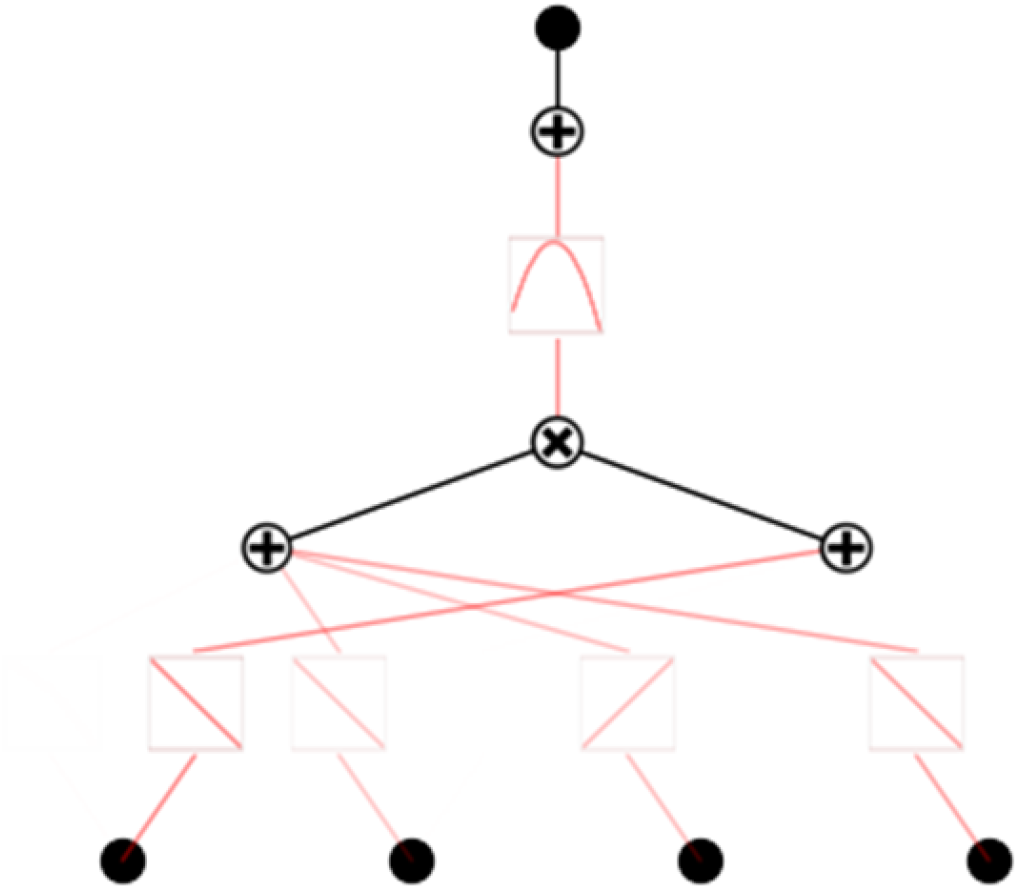
EPIK’s learned intrinsic symbolic graphical representation of the 4-way Multiplicative Interaction scenario.

#### Algorithm 1

Fixed Higher Order XOR Detection

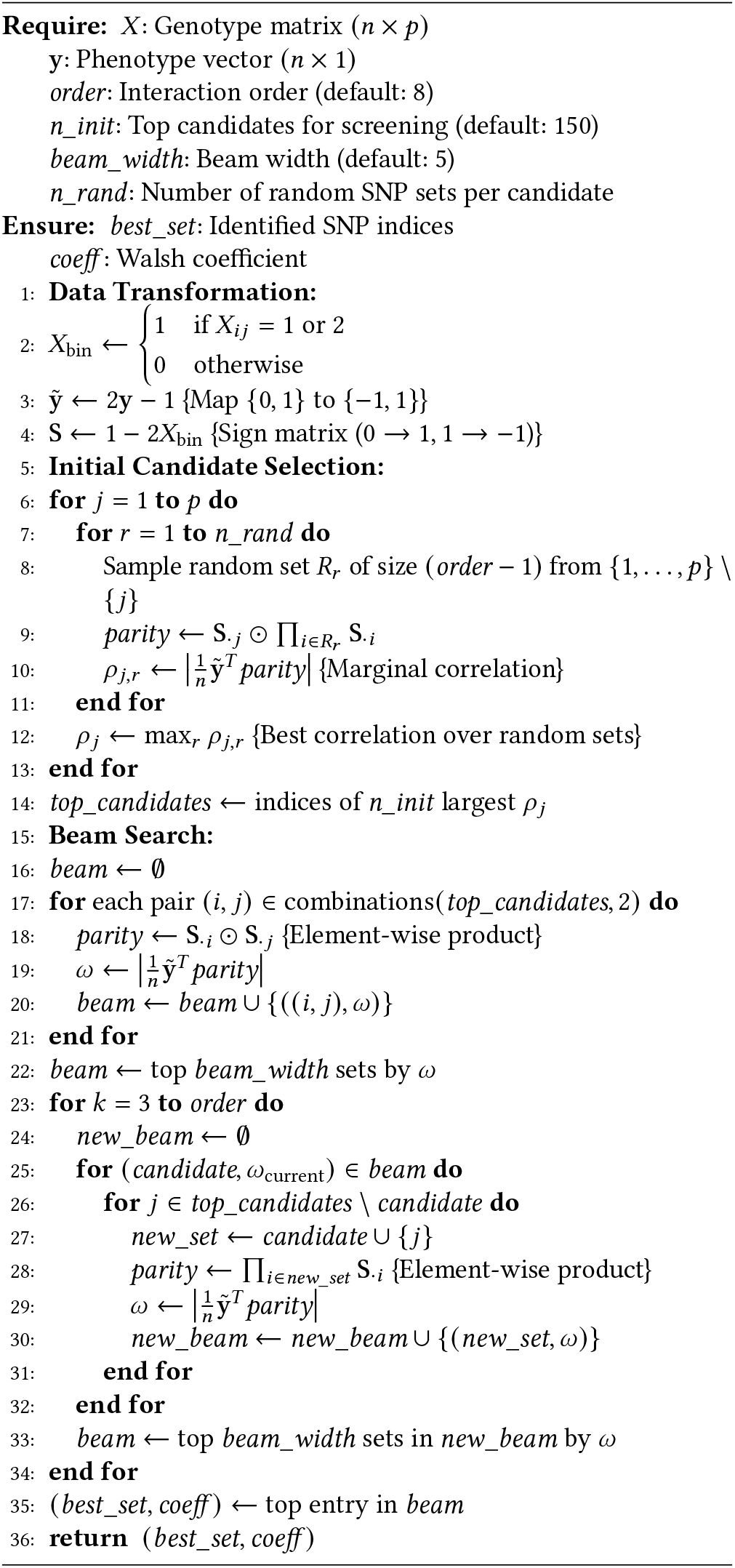

### A.7 Statistical Guarantees discussion of XOR Detection Module and EPIK

For statistical guarantees for the XOR detection module, there aren’t explicit global statistical guarantees as we do not do an exhaustive enumeration. However, we can state that for any of the subsets that are tested, the correlation is an exact estimate for that set. As such, the probability of finding the correct XOR set generally hinges on the actual effect size of the interaction, beam width, the number of initial candidates and their random sets considered, and any traditional quality control procedures, like MAF filtering and LD pruning, keeping the causal SNPs for consideration while reducing feature space. Maximizing beam width and initial candidates/their random sets are what increase the XOR detection module itself for better detecting the interaction as it increases the chance the true set can be found, once found the Walsh coefficient will be much higher than other non-causal sets by nature of XOR’s parity pattern.

For EPIK as a general framework itself, statistical guarantees can be integrated into the framework through splitting the data into a set for selecting candidate SNPs involved in epistasis and using the remaining data to test each candidate SNP set (straifying as needed) with pre-specified generalized linear models dependent on interaction model type and nested likelihood ratio test (LRT) for each with PCs/covariates included as necessary and controlling for FDR. Each test’s p-value for that interaction model’s proposed set can be pooled together and standard Benjamini-Hochberg (BH) at target FDR alpha can be applied. By doing this pooling, it should handle the multiple hypothesis testing involved in epistasis detection.

#### Algorithm 2

Generalized Higher Order XOR Detection

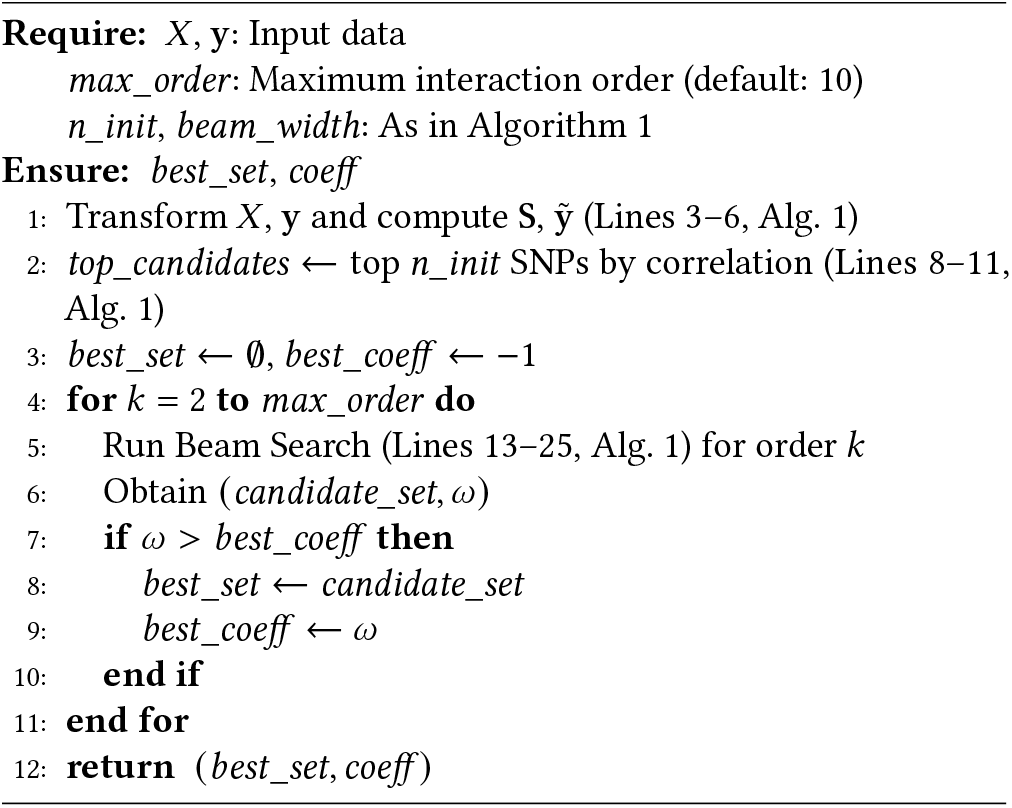

For the higher-arity interactions, we note:

- Candidates increase, resulting in number of tests increasing, causing BH thresholds to decrease and finally resulting in decreased power.
- In general, but especially with higher arity, there should be at least 20-30 samples in this inference set as an initial rule for the framework’s approach.

